# Comparison of online and offline applications of dual-site transcranial alternating current stimulation (tACS) on functional connectivity between pre-supplementary motor area (preSMA) and right inferior frontal gyrus (rIFG) for improving response inhibition

**DOI:** 10.1101/2023.05.03.539327

**Authors:** Hakuei Fujiyama, Alexandra G. Williams, Jane Tan, Oron Levin, Mark R. Hinder

**Author notes:** Corresponding Author: Dr. Hakuei Fujiyama School of Psychology, Murdoch University 90 South Street, Murdoch, WA, 6500, Australia.

## Abstract

**Background:** The efficacy of transcranial alternating current stimulation (tACS) is thought to be brain state-dependent, such that tACS during task performance would be hypothesised to offer greater potential for inducing beneficial electrophysiological changes in the brain and associated behavioural improvement compared to tACS at rest. However, to date, no empirical study has directly tested this postulation.

**Objective:** Here we compared the effects of tACS applied during a stop signal task (online) to the effects of the same tACS protocol applied prior to the task (offline) and a sham control stimulation.

**Methods:** A total of 53 young, healthy adults (32 female; 18-35 yrs) received dual-site beta tACS over the right inferior frontal gyrus (rIFG) and pre-supplementary motor area (preSMA), which are thought to play critical roles in action cancellation, with phase-synchronised stimulation for 15 min with the aim of increasing functional connectivity.

**Results:** EEG connectivity analysis revealed significantly increased task-related functional connectivity following online but not offline tACS. Correlation analyses suggested that an increase in functional connectivity in the beta band at rest following online tACS was associated with an improvement in response inhibition. Interestingly, despite the lack of changes in functional connectivity at the target frequency range following offline tACS, significant improvements in response inhibition were still observed, suggesting offline tACS may still be efficacious in inducing behavioural changes, likely via a post-stimulation early plasticity mechanism.

**Conclusion:** Overall, the results indicate that online and offline dual-site beta tACS are beneficial in improving inhibitory control via distinct underlying mechanisms.

## Introduction

Transcranial alternating current stimulation (tACS) is a form of non-invasive brain stimulation that uses a weak alternating electrical current that is thought to influence the oscillatory timing of brain activity by synchronising endogenous activity with the driving tACS frequency (Herrmann et al., 2013). When applied simultaneously to two cortical sites, tACS can potentially enhance communication, or functional connectivity, between the stimulated brain regions (Polania et al., 2012). Given that various psychiatric illnesses and neurodevelopmental disorders have been associated with problematic functional connectivity, the ability to improve functional connectivity between cortical regions makes tACS particularly appealing as an intervention tool (Tavakoli & Yun, 2017). However, despite the promise that tACS has shown in improving connectivity, the associated effects on behavioural outcomes are inconsistent. It appears whether changes in functional connectivity give rise to associated behavioural change may be mediated by several factors, including task characteristics (e.g., difficulty levels), methodological issues (e.g., electrode placement and timing of the stimulation), and individual variation in cognitive abilities (Brunyé et al., 2019).

Among these issues, one of the important methodological considerations is the state of brain activity during stimulation. As tACS can be applied both during task performance or at rest, it is possible that tACS would interact with distinct electrophysiological processes or plasticity mechanisms, which likely results in different behavioural outcomes (Herrmann et al., 2013). The timing of the stimulation can be classified into ‘online’ or ‘offline’ applications relative to task performance. The online application involves stimulation using a specific frequency relevant to the target behaviour during task performance, which has been associated with dynamic temporal changes in neuron activity (e.g., Veniero et al., 2019). The offline protocol involves stimulation prior to (or post) task performance, which has been associated with stimulation after-effects such as neural plasticity processes; that is, long-term potentiation (LTP) or long-term depression-like (LTD) changes or the possibility of residual temporal changes from active stimulation (Veniero et al., 2019).

Two characteristics of tACS strongly indicate that the online protocol would be superior in inducing changes at both electrophysiological and behavioral levels. Firstly, the effectiveness of tACS is thought to be brain state-dependent: stimulation applied during a task performance that is associated with the applied stimulation frequency would be beneficial (Alagapan et al., 2016; Weinrich et al., 2017). Moreover, as functional connectivity is associated with the timing of dynamic neural activity of network cortical nodes (Fries, 2005, 2015), the effect of tACS can be optimised by the online protocol where it can directly influence temporal dynamics in brain activity (Veniero et al., 2019). While the converging evidence strongly suggests greater neuromodulatory effects with online tACS application, to date, no empirical investigation has directly compared the effects of online and offline tACS applications on functional connectivity. Here, for the first time, we empirically compared the effect of online and offline tACS applications on functional connectivity using electroencephalography (EEG) as well as the associated effects on behaviour in the context of inhibitory control using a double-blind sham-controlled design. Notably, we also considered a sham stimulation condition to control for any placebo-induced changes or changes in performance due to learning effects, independent of the tACS protocol. We specifically chose an inhibitory control task to contrast the behavioural effects of online and offline tACS. This is because (1) deficits in response inhibition have been associated with functional connectivity within the response inhibition network (Tan et al., 2019) and (2) the cortical network that involves the right inferior frontal gyrus (rIFG) and pre-supplementary motor areas (preSMA) together with the subthalamic nucleus (STN) for inhibitory control is well-established (e.g., Aron et al., 2007) and accessible via tACS cortical stimulation. Therefore, the paradigm provides an ideal research framework to explore the effect of tACS applied at different times relative to task performance on electrophysiological activities and behaviour. As it is believed that the effect of tACS is brain-state dependent (Alagapan et al., 2016), it was hypothesised that the extent of improvement in functional connectivity between the preSMA and rIFG, as shown by increased measures of coherence calculated from EEG, would be greater in the online tACS application than the offline application. We further hypothesised that larger improvements in functional connectivity from online stimulation would be associated with greater improvements in inhibitory performance than those produced by offline stimulation. The prediction was based on previous studies utilising dual-site tACS that reported the association between increases in functional connectivity and improved behavioural performance (e.g., Leunissen et al., 2022; Violante et al., 2017). Identifying the specific effects of online and offline protocols when attempting to improve functional connectivity has high clinical importance for developing future interventions for health and disease.

## Material and methods

### Participants

A power analysis was performed for sample size estimation based on data from our previous studies investigating the effect of non-invasive brain stimulation on response inhibition (Fujiyama et al., 2022), motor learning (Fujiyama et al., 2017), and corticospinal excitability (Fujiyama et al., 2014). While the power analysis (alpha = .05, power = .80) indicated that the recommended number of participants to observe a large pre-post change in a group (Cohen’s *d* = 0.8) in response inhibition was 14 participants per group, we recruited at least 17 participants per group to be conservative, which gives a power of .85.

A total of 57 (34 females) healthy volunteers were initially recruited. One participant withdrew from the study and data from two participants were excluded due to technical issues, resulting in a final sample of *N* = 53 (32 females) with an age range of 18-35 years (*M* = 23.07, *SD* = 5.07) who were randomly assigned into the online, offline or sham groups (Table 1). All participants were screened for contraindications for non-invasive brain stimulation, including psychiatric and neuropathological conditions and certain prescription medications (Rossi et al., 2009) and self-reported as having normal or corrected-to-normal vision and hearing. The Edinburgh Handedness Inventory (Oldfield, 1971) was used to assess handedness: only participants with scores above 40 (indicating right-handedness) were included as it has been reported that handedness may cause variations in after effects of non-invasive brain stimulation (e.g., Fitzgerald et al., 2021). Written informed consent was obtained before participation in the study. The study conformed to the Declaration of Helsinki and was approved by the Murdoch University Human Ethics Committee (2016/021).

**Table 1.**

| *Demographic Details of Participants* |  |  |  |
| --- | --- | --- | --- |
|  | Online | Offline | Sham |
|  | *M (SD)* | *M (SD)* | *M (SD)* |
| *N* (Males) | 18 (11) | 18 (11) | 17 (10) |
| Age | 23.1 (4.81) | 24.2 (5.5) | 21.9 (4.9) |
| Handedness<sup>a</sup> | 83.3 (17.8) | 83.9 (17.9) | 86.5 (15.0) |

*Note.* <sup>a</sup> As measured by the Edinburgh Handedness Inventory.

### Apparatus and materials

#### Stop Signal Task (SST)

Inhibitory control was assessed using the SST, which is considered to provide a reliable measure of latent reactive response inhibition (Congdon et al., 2012). PsychoPy (version 1.90.3)(Peirce et al., 2019) was used to configure a two-choice reaction time SST performed on a desktop computer with a monitor refresh rate of 144 Hz. The SST is illustrated in Figure 1A.

**Figure 1.**
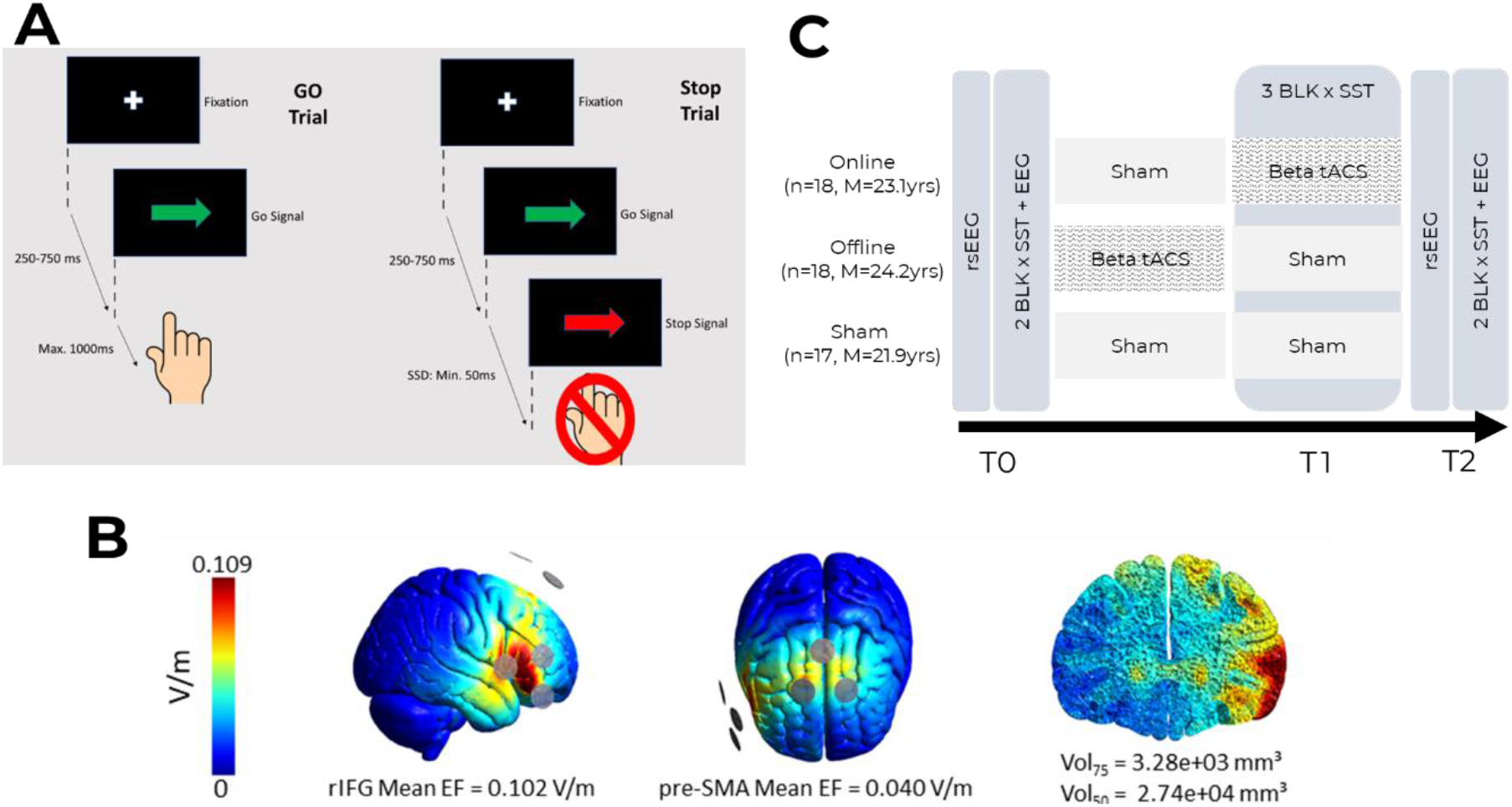
**A.** Stop Signal Task. Participants were instructed to respond to an imperative Go signal (green left- or right-pointing arrow) using their right index finger by pressing the corresponding key on a computer keyboard. Participants kept their index finger on the down arrow key on the keyboard between trials not to bias the direction. During Stop-trials, where the previously-presented green arrow would turn red after a dynamic delay (stop signal delay, SSD) participants had to withhold their response. SSD on the subsequent stop trial was increased, or decreased, by 50 ms following successful or unsuccessful stopping, respectively. **B.** Electrical current flow modelling for the tACS (2 x 1) electrode montage over rIFG and preSMA (Adapted from Tan et al., 2020, with permission of the authors). Vol75 (mesh volume with field strength ≥ 75% of the 99.9^th^ percentile) and Vol50 (mesh volume with field strength ≥ 50% of the 99.9^th^ percentile) provide measures of focality. **C**. Procedural diagram showing differences between online, offline, and sham groups. Electroencephalogram (EEG) measures brain resting-state activity and task-related activity pre and post-tACS period. The online group received tACS concurrently with task performance, while the offline group received tACS prior to task performance at T1. The sham stimulation involves an initial ramp-up of stimulation and ramp-down of stimulation at the end of 15-minute periods.

Each trial began with a white fixation cross in the centre of the black screen, randomly timed to last between 250 and 750 ms (drawn from a uniform distribution), which was then followed by an imperative Go signal in the form of a green arrow either pointing left or right for a duration maximum of 1000ms. Participants were instructed to press the corresponding left or right arrow key on a qwerty keyboard with their right index finger as fast as possible after the presentation of the Go signal. Participants were explicitly instructed to keep their finger on the down arrow key on the keyboard between trials not to bias the direction. The time between the onset of the Go imperative signal and registration of the finger pressing down was the Go Reaction Time (Go RT). Visual feedback (i.e., displaying the Go RT in ms) was provided after each successful trial for 1000 ms to motivate participants to respond as quickly as possible. When participants failed to respond to the Go signal or pressed the wrong button, “missed” was displayed on the screen in lieu of RT feedback.

In 25% of trials, the green Go signal was replaced SSD ms after the presentation by a red arrow facing the same direction that served as a Stop signal, which necessitated that participants withhold their selected (initiated) response. SSD was manipulated in a staircase fashion with the initial SSD set at 200 ms; SSD increased or decreased by 50 ms in the subsequent stop trial depending on successful or failed stopping, respectively (Allen et al., 2018; Aron & Poldrack, 2006). In this manner, we aimed to achieve approximately 50% stopping success. In Stop-trials, a successful stop was followed by the visual feedback "good", and an unsuccessful stop was followed by “please try to stop”. Trials were presented in blocks of 80 trials (60 Go trials, 20 Stop trials), and mean Go RT and stop success rate (%) were presented as a summary feedback screen at the completion of each block.

### Transcranial alternating current stimulation (tACS)

Dual-site tACS was applied using a neuroConn DC-STIMULATOR MC machine (NeuroConn, Ilmenau, Germany). Round rubber electrodes (2cm diameter) were placed over scalp sites targeted at the rIFG and the preSMA and held in place with conductance paste. Alternating currents were applied at the beta frequency of 20 Hz, with a current intensity of 1 mA peak-to-peak amplitude and with zero DC offset to both stimulation sites at 0° phase lag, i.e., in-phase. Electrical impedance was kept under 50 kΩ (Spitoni et al., 2013). For online and offline groups, tACS was applied for 15 mins, inclusive of a 30-second ramp-up and ramp-down of current intensity. Sham stimulation consisted of the initial 30-second ramp-up period and post 15 minutes ramp-down period of current only.

The electrode placement sites were identified using the international 10-20 system (Herwig et al., 2003). The rIFG was located at the intersection between Fz to T4 and Cz to F8, and the preSMA was located at Fz (Herwig et al., 2003). A 2 x 1 montage, consisting of two active electrodes and one reference electrode, was applied to each location with 2 cm between the centre of the electrodes and the estimated target location (Figure 1B) It is recommended that current flow modelling is used to confirm focal current distribution to sites of interest (Karabanov et al., 2019). Electrical field modelling (Tan et al., 2020; Figure 1B) indicated that the 2 x 1 montage produces adequate focal current distribution to the rIFG and preSMA. The tACS machine was pre-programmed by a research associate with codes representing the three conditions to ensure the blinding of both participants and the researcher.

### Electroencephalogram (EEG)

EEG was recorded using HydroCel Geodesic Sensor Nets (Magstim EGI, Eugene, OR) that have 128 scalp electrodes with the HydroCel GSN 128 1.0 montage and Net Station (4.5.6) software. EEG was amplified with Net Amps 300 amplifier, low and high pass filtered (0.1–500 Hz), and digitally converted at a sampling rate of 1000 Hz. Signals were referenced to the Cz referencing during recording, and electrical impedance was kept below 50 kΩ as per the manufacturer’s recommendation (Magstim EGI, Eugene, OR).

#### Pre-processing

MATLAB (Mathworks, R2018b) and the EEGLAB toolbox (Delorme & Makeig, 2004) was used to pre-process the resting-state EEG from pre- and post-tACS. Data were down-sampled to 500 Hz and run through Hamming windowed-sinc finite impulse response (FIR) filters (high-passed at 1 Hz and notch filtered at 50, 100, and 150 Hz) to remove slow drifts and electrical noise. Bad channels and segments were manually removed via visual inspection. The excluded channels were interpolated, and all signals were re-referenced to the average. Finally, an independent component analysis (EEGLAB toolbox *runica()*) option) was used to remove ocular, muscular and vascular artefacts.

The processed data was then run through customised Matlab scripts and the Brainstorm toolbox (Tadel et al., 2011) to determine connectivity between the two stimulation sites.

### Control measures

#### Sleep, caffeine, alcohol, cigarette questionnaire

To account for possible confounding effects (e.g., sleep quality and quantity, caffeine, alcohol, and cigarette consumption) on the effects of tACS, a questionnaire was administered to monitor inter-group variability. This consisted of five independent items with scales. Item 1 required participants to rate their sleep quality on a scale from 1 to 10, with 1 representing very poor quality. Items 2 to 5 were the number of hours slept during the previous night relative to the day of the testing, as well as the units of caffeine and alcohol consumed and the number of cigarettes smoked in the 12 hours before testing.

#### Non-invasive Brain Stimulation Sensation (NiBS) Questionnaire

To assess the efficacy of blinding and detect any adverse effects, participants completed a NiBS questionnaire that asked about any sensations they might have experienced during stimulation. There were 12 items that included sensations previously associated with NiBS, such as “itching”, “headache’, and “tingling” (Woods et al., 2016). Participants rated these sensations on a 5-point scale, which ranked options from “nothing” to very strong”, and one item questioning if the sensations influenced task performance rated on a 5-point scale, with response options ranging from "a little" to "very much". Item scores were combined for a range of scores from 13-65. There were 3 items reporting sensation duration, with a 3-point scale with the options for “start”, “middle”, and “end” of stimulation session.

### Procedure

None of the participants had ever received any form of non-invasive brain stimulation prior to participating in the present study. The (blinded) experimenter (AGW) used standardised instruction across the groups to minimise the variation in any placebo effect, as any expectations in a particular group may influence the outcome of the stimulation (Peerdeman et al., 2015). Figure 1C illustrates the experimental flow. Each participant completed a single experimental session which began with a three-minute eyes-open resting-state EEG (rsEEG) recording (T0). After an initial 16 practice trials of SST to ensure task comprehension, two blocks (80 trials each) of SST were then performed (T0) during the continuous acquisition of EEG, i.e., during the entire block, with a short break in-between the blocks. To encourage participants to maximise their task performance, they manually recorded their mean Go RT and performance scores (number of successful stops) on a score sheet after each SST block.

Over the next ∼30 minutes, in-phase beta tACS or sham stimulation were applied over the rIFG and preSMA before or during SST: The online group started with sham stimulation at rest, followed by in-phase beta tACS during the performance of SST (T1), while the offline group received in-phase beta tACS first at rest, followed by sham stimulation during the performance of SST (T1). The sham group received sham stimulation both at rest and during the performance of SST (T1). This design permits full blinding for the experimenters both during the experiments (AJW) and during data analysis (HF, JT) in regards to online/offline/sham, with sham condition permitting any changes in SST performance and EEG to be assessed in the absence of real tACS. Moreover, the design permits comparable assessment time points across groups, which were instrumental for accurately comparing the effects of online and offline tACS applications.

The sham stimulation condition involved an initial ramp-up and a final ramp-down of stimulation during both 15-minute periods (Fig 1 C). In all groups, participants rested during the first 15 min and completed three training blocks of SST (T1) in the subsequent 15 min period.

On completion of three SST blocks, three-minute eyes open rsEEG was again recorded (T2), followed by two blocks of SST performance (T2). At the end of each session, participants completed the NiBS questionnaire. Each session lasted for approximately 2.5 to 3 hours.

### Data processing and statistical analysis

#### Control measures

Items in the sleep, caffeine, alcohol, and cigarette questionnaire and NiBS sensations questionnaire were individually analysed using a Kruskal-Wallis test with the between-subject independent variable (IV) of GROUP (online, offline, sham). For the NiBS sensations questionnaire comparing sensations between groups, a one-way between-groups ANOVAs with as GROUP (online, offline, sham) as IV was used.

#### Behavioural measures

Go RT is the response time (i.e., mechanical button press) to the choice imperative stimuli without the subsequent stop signal presentation excluding incorrect responses, and was used to assess response speed in the Go component of the task. Stop Signal Reaction Time (SSRT) was used to assess reactive inhibitory control (i.e., action cancellation when the unexpected stop signal appears) and was calculated using the integration method (Verbruggen et al., 2019), which is less affected by stopping success rates that deviate from 50% (Band et al., 2003). Specifically, the mean SSD was calculated for each participant at each time point separately. Initially all go trials including go trials with a choice error and go trials with a premature response, were initially included in SSRT calculation (Verbruggen et al., 2019). Since SSRT estimation relies on the assumption of an independent race between the go and stop processes (e.g., Verbruggen & Logan, 2009), for each participant and time, we excluded conditions if the mean RT on unsuccessful stop trials was greater than the mean Go RT in that same condition (Verbruggen et al., 2019). Furthermore, for each participant and for each time point, conditions with stop accuracy < 0.25 or > 0.75 were excluded, as suggested by Congdon and colleagues (2012) and Verburggen et al. (2019). Go RT for correct responses in the corresponding trials was arranged in ascending order and the Go RT corresponding to the stop success rate (i.e., 1-percent successful inhibition) was identified. For example, if the stop success was 55%, the 55^th^ percentile Go reaction time was identified as the quantile reaction time and the mean SSD was subtracted from the quantile reaction time to estimate SSRT. We also excluded SSRTs less than 50 ms from the statistical analyses (Congdon et al., 2012). For Go RT analysis (but not for the calculations of SSRT), any trials with less than 150 ms of Go RT were excluded as they were deemed to be too fast.

#### EEG measures

Source localisation was performed with depth-weighted linear L2-minimum norm estimates (Baillet et al., 2001; Gramfort et al., 2014), through the Brainstorm toolbox (Tadel et al., 2011). The forward model was constructed with the ICBM152 template brain (Fonov et al., 2009) and the Symmetric Boundary Element Method (BEM) implemented in the OpenMEEG software (Gramfort et al., 2010; Kybic et al., 2005). The AAL2 volume atlas was used for cortical and subcortical parcellation (Rolls et al., 2015; 103 regions), with additional parcellations for the preSMA, SMA proper, and the subthalamic nucleus from the ATAG atlas (Keuken et al., 2014).

Analytical signals from the resultant source localised data were convolved with complex Morlet wavelets in 1-Hz increments for frequencies between 8 and 45 Hz. The wavelet lengths ranged from 4 to 10 cycles in 38 linearly-spaced steps, with the wavelet length increasing with the wavelet frequency for the dynamic adjustment of the balance between temporal and frequency precision (Cohen, 2014). To minimize the effects of edge artifacts, the resultant analytic signals were analysed in time windows of 400 to 1600 ms in each epoch for resting-state data and -200 to 1000 ms during successful stop trials for task-related data.

The imaginary component of coherency (ImCoh), an index of the consistency of phase angle differences (phase lag) between signals (Nolte et al., 2004), was used to assess changes in phase-coupling that were induced by tACS. For channels *i* and *j*, with the complex Fourier transforms *x*_*i*_(*f*) and *x*_*j*_(*f*) of their time series data, ImCoh at frequency *f* is given as

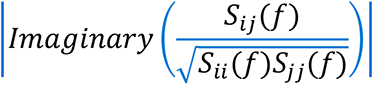

where the cross-spectral density *S*_*ii*_(*f*) is derived from the complex conjugation of *x*_*i*_(*f*) and *x*_*j*_(*f*). The coherency between the channel signals is obtained by normalising the cross-spectral density by the square root of the signals’ spectral power (*S*_*ii*_(*f*) and *S*_*jj*_(*f*)). The result is a complex number from which the imaginary component, ImCoh, is extracted.

ImCoh has been shown to be insensitive to the effects of volume conduction, and its usage thus reduces the likelihood of mistakenly taking spurious connectivity for real functional coupling (Nolte et al., 2004). An ImCoh value of 1 denotes perfect phase-coupling between signals, while a value of zero indicates that the phase angle differences between the signals are completely random. The ImCoh was calculated for each inter-site channel-pair (F4-Fz, F6-Fz, F4-AFz, and F6-AFz) and subsequently averaged across the channel-pairs to result in the mean ImCoh estimate between the rIFG and preSMA.

The estimates of ImCoh were computed at the trial-level, in sliding time windows with frequency-varying lengths of 3-8 cycles in 38 linearly-spaced steps, at 20 ms intervals within each epoch. These estimates were subsequently averaged within and across epochs to obtain an ImCoh estimate for each participant at each condition.

#### Statistical Analysis

A generalised linear mixed model (GLMM) was constructed to analyse Go RT and ImCoh data, while a linear mixed model (LMM) was used to analyse SSRT; GROUP (online, offline, sham) and TIME (T0, T1, T2 for Go RT and SSRT; T0, T2 for ImCoh) were fixed factors with by-subject intercept as a random effect. As the Go RT and ImCoh data were continuous, positive, right-skewed, and/or where variance was near-constant on the log scale, GLMMs were used, applying a gamma distribution with a log link function (Lo & Andrews, 2015). For the GLMMs analysing Go RT and ImCoh, data at the single trial level were used, i.e., Go RT for each go trial and average ImCoh for each resting-state epoch and successful stop trial. A LMM was used to analyse SSRT, obtaining a mean SSRT for each time point for each individual participant (as SSRT cannot be calculated at the single trial level) – normality of these mean data is assumed across individuals/time points and hence a LMM is conducted. For ImCoh data, we investigated stimulation-specific effects focusing on 20 Hz ImCoh as well as on broader frequency ranges, including alpha (8–12 Hz), beta (13–30 Hz) beta, and gamma (30–45 Hz) bands (Schmiedt-Fehr et al., 2016). Null hypothesis significance testing for main and interaction effects was conducted using Wald Chi-Squared tests for GLMM and *F*-tests for LMM with significant main and interaction effects were further investigated with custom Bonferroni-corrected contrasts. We also performed a series of Pearson’s product-moment correlations to explore the relationship between the extent of changes in functional connectivity indexed by ImCoh and the extent of behavioural changes following the application of tACS. For this purpose, change (Δ) of T2/T0 SSRT and change (Δ) of T2/T0 ImCoh were calculated for each frequency band as well as for 20 Hz. Statistical significance was set at *p* < 0.05. For the correlation analyses, given the exploratory nature of these analyses and the substantial number of comparisons, we opted not to apply corrections for multiple comparisons (Antonenko et al., 2019; Fujiyama, Van Soom, Rens, Cuypers, et al., 2016; Fujiyama, Van Soom, Rens, Gooijers, et al., 2016). As such, these correlations need to be interpreted with caution.

All statistical analyses and visual illustrations of the results were performed using the R statistical package, version 4.1.3 (R Development Core Team, 2021) with an integrated environment, RStudio version 2023.03.0.386 (RStudio Team, 2020) for x86_64-w64-mingw32/x64 running under Windows 10 version 20H2 using pwr 1.3.0 (Champely, 2020) for power analysis, car version 3.0-8 (Fox et al., 2020) for ANOVAs, lme4 1.1-31 (Kuznetsova et al., 2017) for GLMM and LMM fitting, emmeans version 1.8.4.1 (Lenth, 2020) for post-hoc comparisons, and ggplot2 version 3.2.1 (Wickham et al., 2020).

## Results

### Control measures

Descriptive statistics for control measures (i.e., from the NiBS sensation questionnaire and the sleep, caffeine, alcohol, and cigarette questionnaire) were reported in Table 2.

**Table 2.** Descriptive statistics for the control measures

| Measure | Online | Offline | Sham |  |
| --- | --- | --- | --- | --- |
|  | *M (SD)* | *M (SD)* | *M (SD)* | *p*-value |
| NiBS sensation (0-4) | 1.44 (0.19) | 1.45 (0.19) | 1.40 (0.18) | 0.758 |
| Sleep Quality (1-10) | 7.71 (0.27) | 6.65 (0.50) | 6.71 (0.44) | 0.318 |
| Sleep Quantity (hour) | 7.56 (0.39) | 6.79 (0.37) | 6.91 (0.33) | 0.459 |
| Caffeine (unit) | 0.65 (0.21) | 1.35 (0.36) | 0.88 (0.32) | 0.278 |
| Alcohol (unit) | 0.35 (0.21) | 0.18 (0.17) | 0.00 (0.00) | 0.166 |
| Cigarette | 0.18 (0.18) | 0.00 (0.00) | 0.00 (0.00) | 0.601 |

*Note.* The *p*-values represent the results of the independent-sample Kruskal-Wallis tests, except for the NiBS sensation questionnaire, for which a one-way between-groups ANOVA was used.

A one-way between-group ANOVA (GROUP: online, offline, and sham) was used to investigate the efficacy of blinding by comparing the scores on the NiBS sensation questionnaire, which indexes the perceived sensations during tACS. No significant differences were found between the groups, *F*(2, 49) = 0.278, *p* = .758, *η_p_*² = .012, indicating successful blinding between the groups. Similarly, for the sleep, caffeine, alcohol, and cigarette questionnaire, there were no statistically significant differences between the groups (*p*s > 0.166). As control measures did not show differences between the groups, any effect in the main outcome measure is unlikely to be attributable to these factors.

### Functional connectivity measure: imaginary component of coherency (ImCoh)

We investigated whether the timing of tACS relative to the task performance changes the functional connectivity between preSMA and rIFG in the source space both for resting-state and task-related EEG data. For this, we considered the stimulation-specific effects focusing on ImCoh at 20 Hz as well as in broader frequency ranges, including alpha (8–12 Hz), beta (13–30 Hz) beta, and gamma (30– 45 Hz) bands.

#### Stimulation-specific frequency effect

##### Resting-state ImCoh

For resting state ImCoh at 20 Hz, there were no significant main effects or interactions involving GROUP (online, offline, sham) and TIME (T0, T2), χ*^2^*s < 1.91, *p*s > .150.

##### Task-related ImCoh

ImCoh data from the successful stop trials to investigate the *functional* interactions between the rIFG and preSMA were also analysed to investigate whether the timing of tACS affects functional connectivity *during* the task performance. We found a significant interaction between GROUP and TIME, χ*^2^* (*N* = 53) = 6.96, *p* = .031, which revealed that the online group showed a significant increase (4.98%) in 20 Hz ImCoh value from T0 to T2, *z* = -2.84, *p* = .005, *d* = -0.21 (Figure 2), whereas both the offline (1.44% reduction from T0 to T2) and sham group (0.66% increase from T0 to T2) did not show a statistically significant change in ImCoh from T0 to T2, *p* = .404, *d* = 0.06 (offline group) and *p* = .703, *d* = -0.03 (sham group).

**Figure 2.**
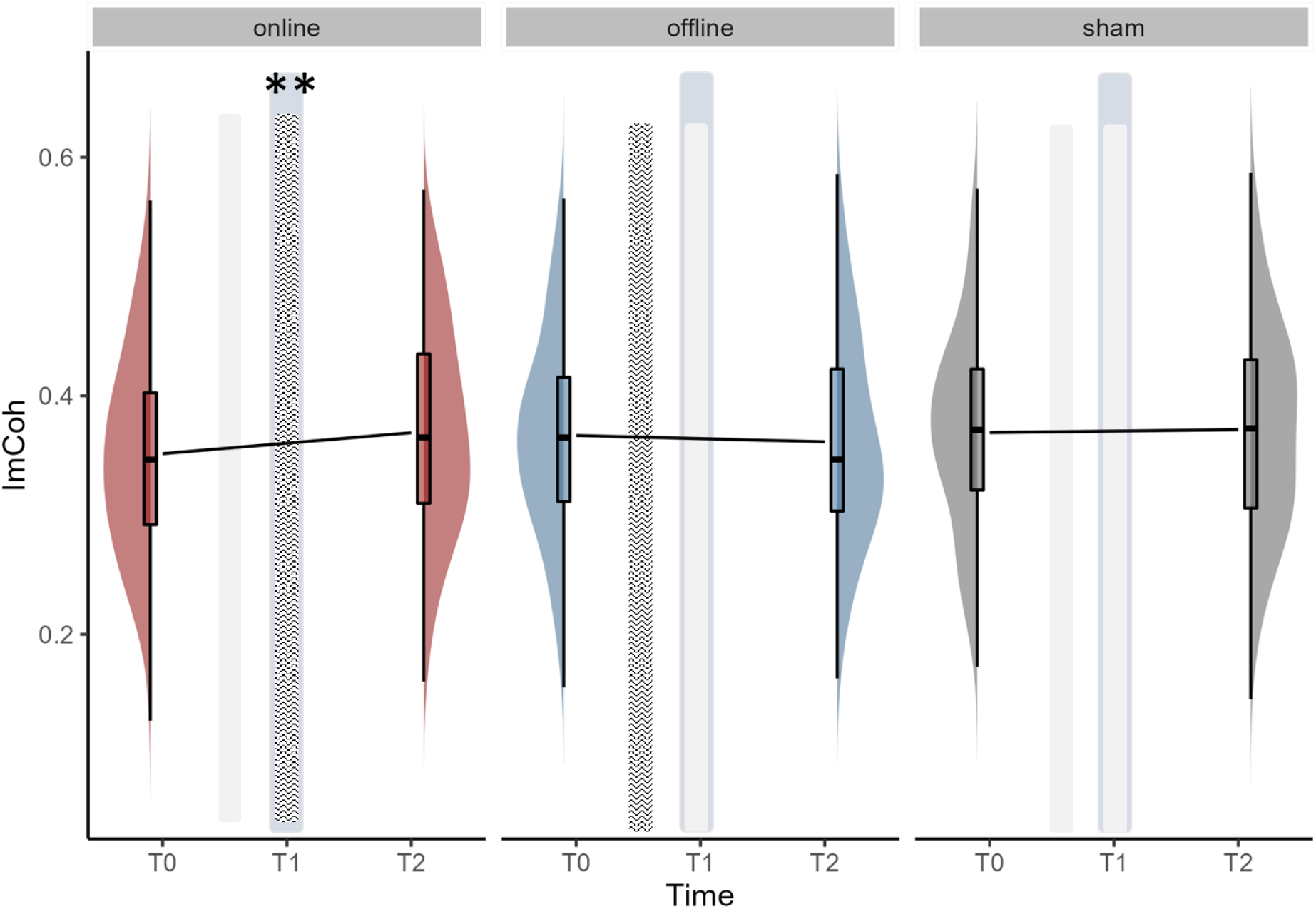
Pre- to Post-tACS changes in task-related imaginary component of coherency e (ImCoh) at 20 Hz during successful stop trials for each group. The shaded vertical bars represent the timing of tACS application, while the light grey vertical bars represent the timing of sham stimulation. The blue-grey vertical bar represents the timing of 3 blocks of SST performance. For the boxplots, the horizontal line within the box plot represents the median. The top and bottom lines of the boxplots represent the upper and lower quartile, respectively. The vertical lines of the box plots represent the distribution showing the minimum and maximum values. The data points outside of the whiskers represent data that are > 1.5 quartiles. Asterisk (∗) denotes a significant effect over time. ** *p* < .01.

#### Broader frequency ranges

##### Resting-state imaginary component of coherency (ImCoh)

We observed a significant main effect of FREQUENCY and significant interactions of GROUP x FREQUENCY and TIME x FREQUENCY, which were mediated by a significant interaction between GROUP, TIME, and FREQUENCY, χ*^2^*(*N* = 53) = 14.16, *p* = .007. As illustrated in Figure 3A, a significant increase (0.86%) in ImCoh was observed at the beta range in the online group, *z* = -4.09, *p* < .001, *d* = -0.10, whereas ImCoh significantly decreased (1.12%) from T0 to T2 in the alpha range, *z* = 2.88, *p* = .004, *d* = 0.13 in the online group. Other groups did not show any statistically significant changes from T0 to T2, *z*s < 1.17, *p*s > .242, |*d*s| > 0.041. Other main effects and interactions were not significant, χ*^2^*s < 5.06, *p*s > .080.

**Figure 3.**
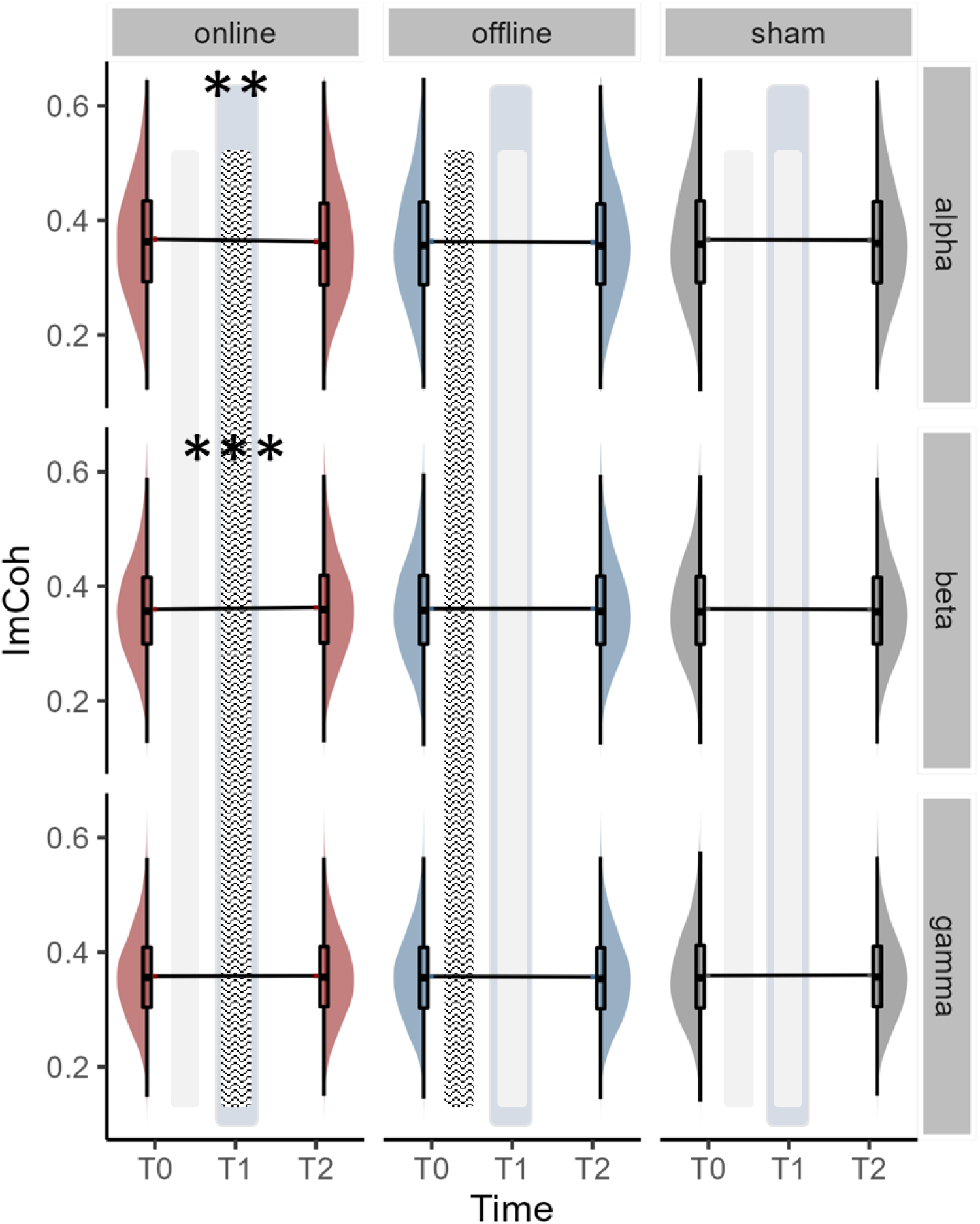
Pre- to Post-tACS changes in *resting state* imaginary component of coherency (ImCoh) across alpha, beta and gamma frequency bands. The shaded vertical bars represent the timing of tACS application, while the light grey vertical bars represent the timing of sham stimulation. The blue-grey vertical bar represents the timing of 3 blocks of SST performance. For the boxplots, the horizontal line within the box plot represents the median. The top and bottom lines of the boxplots represent the upper and lower quartile, respectively. The vertical lines of the boxplots represent the distribution showing the minimum and maximum values, excluding outliers at the tip. The data points outside of the whiskers represent data that are > 1.5 than quartile. Asterisk (∗) denotes a significant effect over time. *** *p* < .001, ** *p* < .01.

##### Task-related imaginary component of coherency (ImCoh)

ImCoh data during the successful stop trials were also analysed to investigate the timing of tACS application on the functional interaction during the task performance. All of the main effects and interactions were statistically significant, χ*^2^*s < 16.71, *p*s > .001, except for the main effect of GROUP, χ*^2^* (*N* = 53) = 4.26, *p* = .119. These effects were best interpreted by a significant 3-way interaction between GROUP, TIME, and FREQUENCY, χ*^2^* (*N* = 53) = 53.30, *p* < .001. The 3-way interaction, together with reference to Figure 4, indicates that the online beta tACS application was beneficial in improving functional connectivity in the beta range during a successful stop trial, exhibiting a greater extent of increase in ImCoh (3.31% increase) from T0 to T2, compared to the changes in the offline (1.21% decrease), *z* = -7.98, *p* < .001, *d* = - 0.13, and sham groups (1.31% increase), *z* = -3.62, *p* < .001, *d* = -0.13. Interestingly, we observed that in the offline group, ImCoh values significantly decreased from T0 to T2 in beta, *z* = 2.95, *p* = .003, *d* = 0.05, and gamma bands (2.92% decrease), *z* = 6.46, *p* < .001, *d* = 0.13, while a significant increase in ImCoh was observed in the alpha band (3.06% increase), *z* = -3.99, *p* < .001, *d* = -0.14. There were no significant changes from T0 to T2 in any of the frequency bands for the sham group.

**Figure 4.**
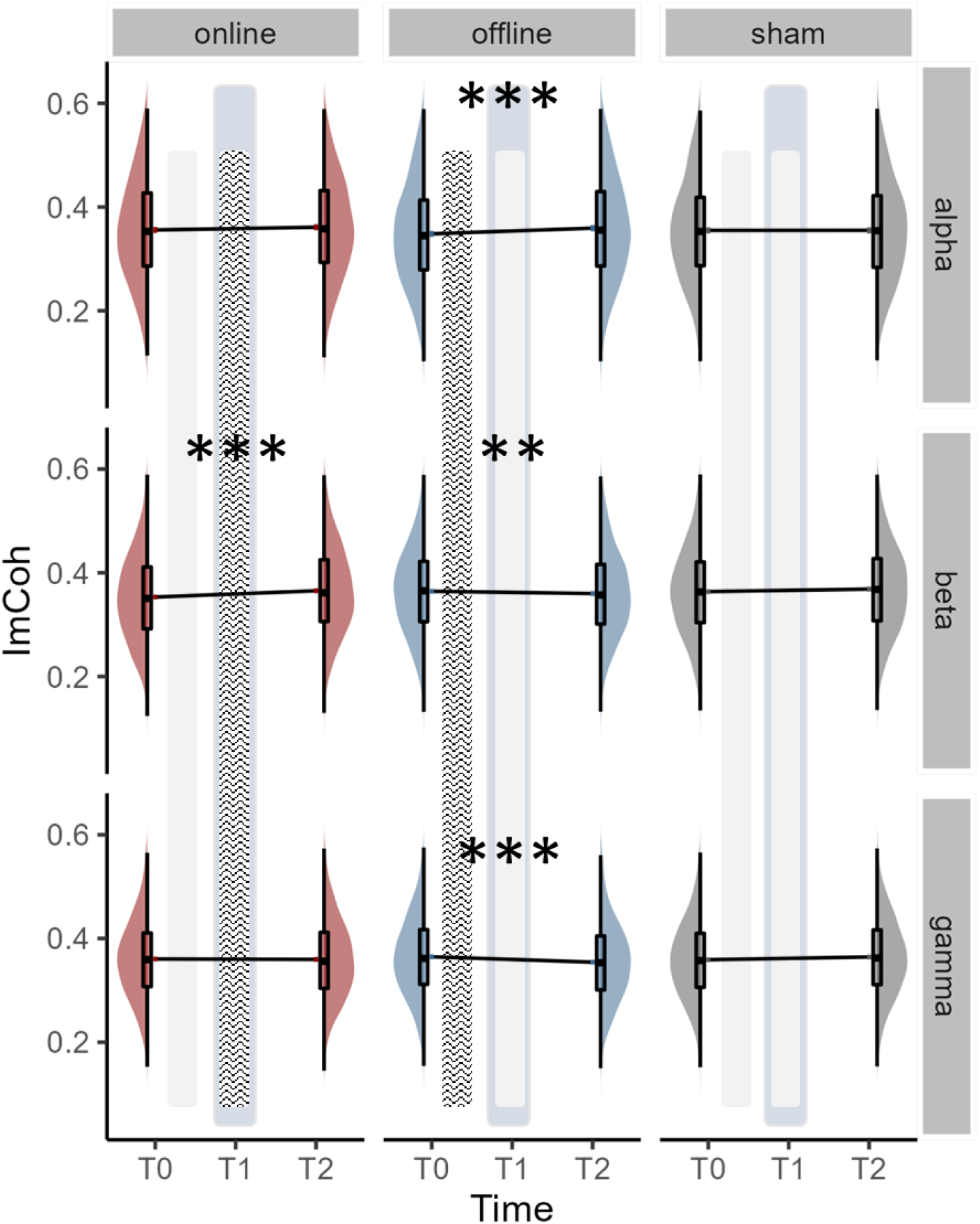
Pre- to Post-tACS changes in *task-related* imaginary component of coherency (ImCoh) across alpha, beta and gamma frequency bands during successful stop trial for each group. The shaded vertical bars represent the timing of tACS application, while the light grey vertical bars represent the timing of sham stimulation. The blue-grey vertical bar represents the timing of 3 blocks of SST performance. For the boxplots, the horizontal line within the box plot represents the median. The top and bottom lines of the boxplots represent the upper and lower quartile, respectively. The vertical lines of the boxplots represent the distribution showing the minimum and maximum values, excluding outliers at the tip. The data points outside of the whiskers represent data that are > 1.5 than quartile. Asterisk (∗) denotes a significant effect over time. *** *p* < .001, ** *p* < .01.

### Behavioural measures

*Go Reaction Time (Go RT).* A significant main effect of GROUP, χ*^2^* (1, *N* = 53) = 7.39, *p* = .025, was observed, although post-hoc comparisons indicate that slower Go RT in the sham group (*M* = 446.17 ± 1.80 ms) compared to the online (*M* = 434.22 ± 1.66 ms) or offline (*M* = 435.79 ± 1.83 ms) groups did not reach the conventional statistical significance level (*z*s < - 2.35, *p*s > .056, |*d*s| < -0.181). The main effect of TIME, χ*^2^* (1, *N* = 53) = 227.6, *p* <. 001, and the associated post-hoc contrasts indicated the presence of a general learning effect: Go RT was faster at T1 (i.e., during the performance of SST, *M* = 434.97 ± 1.50 ms) and T2 (i.e., post, *M* = 433.86 ± 1.83 ms) than T0 (i.e., pre, *M* = 435.79 ± 2.05 ms), *z*s > 13.37, *p*s < .001, |*d*s| > 0.223 (Figure 5). The interaction between GROUP and TIME was not significant, χ*^2^* (1, *N* = 53) = 6.86, *p* = .143, indicating that all the groups showed similar changes in Go RT across time points. Notably, the improvement in Go RT likely mean that participant did not adapt proactive slowing strategies to try and improve stopping success .

**Figure 5.**
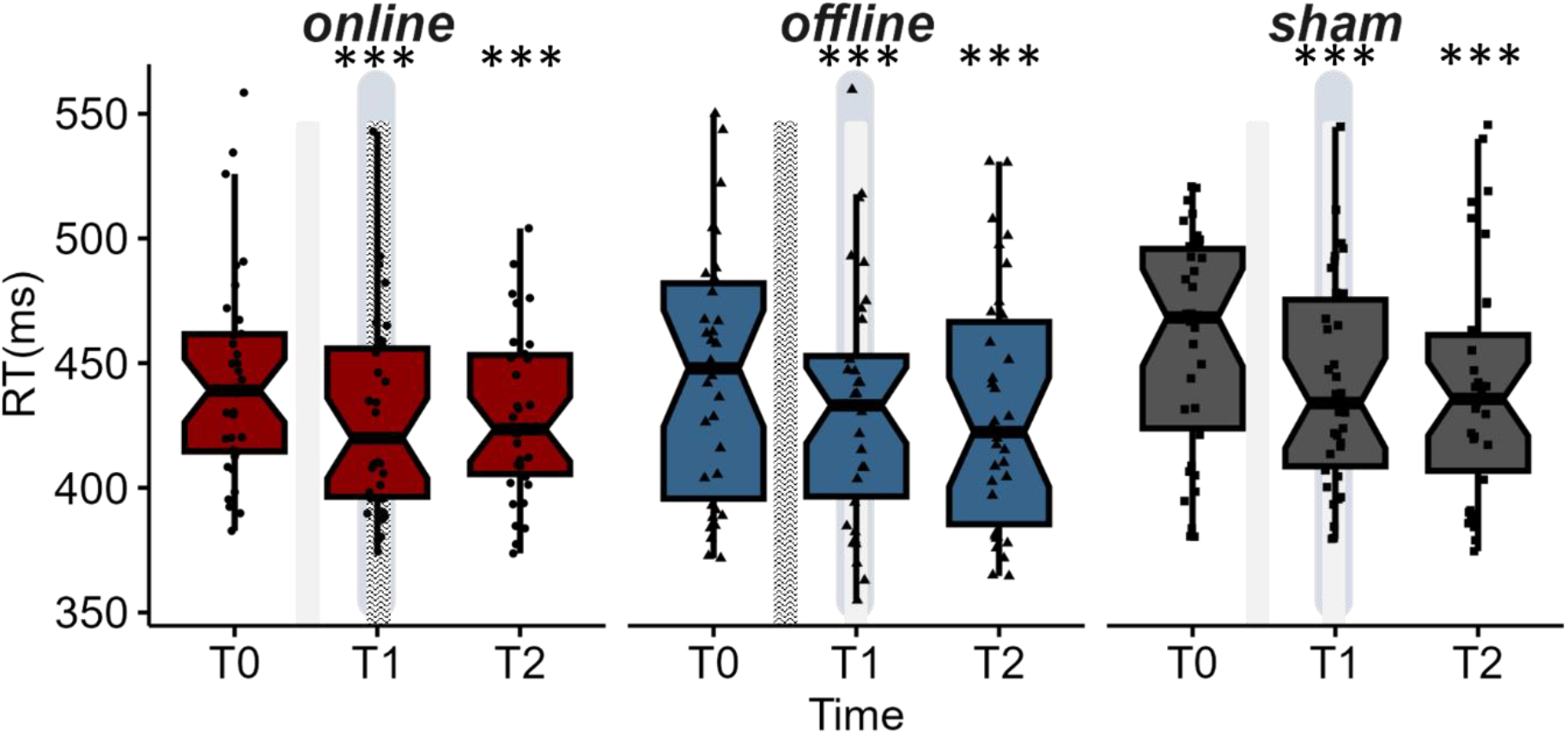
Reaction Time in milliseconds across groups and time points. The shaded vertical bars represent the timing of tACS, while the light grey vertical bars represent the timing of sham stimulation. The blue-grey vertical bar represents the timing of 3 blocks of SST performance. For the boxplots, the horizontal line within the box plot represents the median. The top and the bottom lines represent the upper and lower quartile, respectively. The vertical lines represent the distribution showing the minimum and maximum values, excluding outliers at the tip. The data points outside of the whiskers represent data that are > 1.5 than quartile. The notch displays a confidence interval around the median based on the median ± 1.58*interquartile range/sqrt(n). Asterisk (***) denotes a significant effect over time. *** *p* < .001.

#### Stop signal reaction time (SSRT)

We observed a significant main effect of TIME, *F*(2, 537.41) = 4.97, *p* = .007, which was mediated by the higher-order interaction between GROUP and TIME, *F*(4, 537.31) = 2.99, *p* = .018. As shown in Figure 6, in the offline group, SSRT significantly improved (shorter SSRT) 5.24% from T0 to T1, *p* = .033, *d* = 0.44, and the improvement remained significant at T2 relative to T0 (5.08% reduction in SSRT), *t* = 3.39, *p* = .002, *d* = 0.64. In contrast, SSRT changes in both the online and sham groups were not statistically significant, *t* < 2.34, *p*s > .060, *d*s < 0.42.

**Figure 6.**
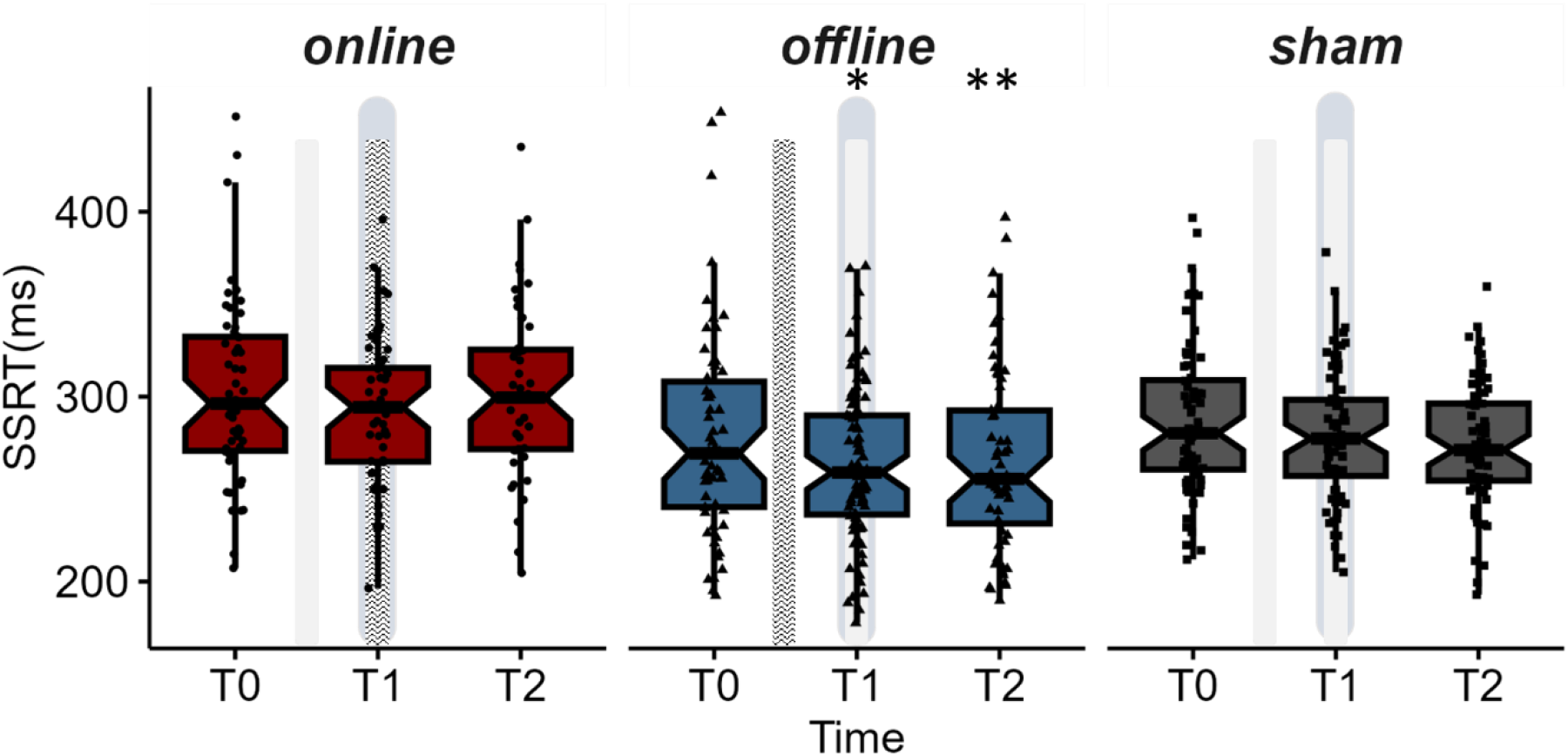

Stop signal reaction time (SSRT) in milliseconds across time points for online, offline, and sham groups. The shaded vertical bars represent the timing of tACS application, while the light grey vertical bars represent the timing of sham stimulation. The blue-grey vertical bar represents the timing of 3 blocks of SST performance. For the boxplots, the horizontal line within the box plot represents the median. The top and the bottom lines represent the upper and lower quartile, respectively. The vertical lines represent the distribution showing the minimum and maximum values, excluding outliers at the tip. The data points outside of the whiskers represent data that are > 1.5 than quartile. The notch displays a confidence interval around the median based on the median ± 1.58*interquartile range/sqrt(n). Asterisk (∗) denotes a significant effect over time. ** *p* < .01, * *p* < .05.

### Neural correlates of response inhibition

Pearson’s product-moment correlation coefficients (*r*) were obtained to explore the relationship between the extent of changes in functional connectivity and changes in behavioural performance by correlating ΔImCoh (i.e., ImCoh at T2/ImCoh at T0) value at rest and during the successful stop trials in each frequency category separately with ΔSSRT (i.e., SSRT at T2/SSRT at T0) for stimulation group separately at 20 Hz and in each frequency band (Table 3). As shown in Figure 7, we observed a medium correlation between resting state ΔImCoh in the beta band (13–30 Hz) and ΔSSRT in the online group, *r* = -0.49, *p* = .038, suggesting that a greater improvement in resting state ImCoh in the beta band was associated with a greater reduction in SSRT, i.e., improvement in response inhibition. Although it did not reach the conventional significance level, i.e., *p* < .05, task-related ΔImCoh in the beta band *during successful stops* showed a similar negative correlation between ΔImCoh and ΔSSRT with a small effect size (*r* = -.204) for the online group. In the sham group, while the correlation coefficient failed to reach the conventional significance level of .05, the effect size representing the relationship between ΔImCoh in the alpha band *during successful stop* and ΔSSRT was close to medium (*r* = -0.468, *p* = .068). Lastly, for the offline group, correlation coefficients were all small and non-significant.

**Figure 7.**
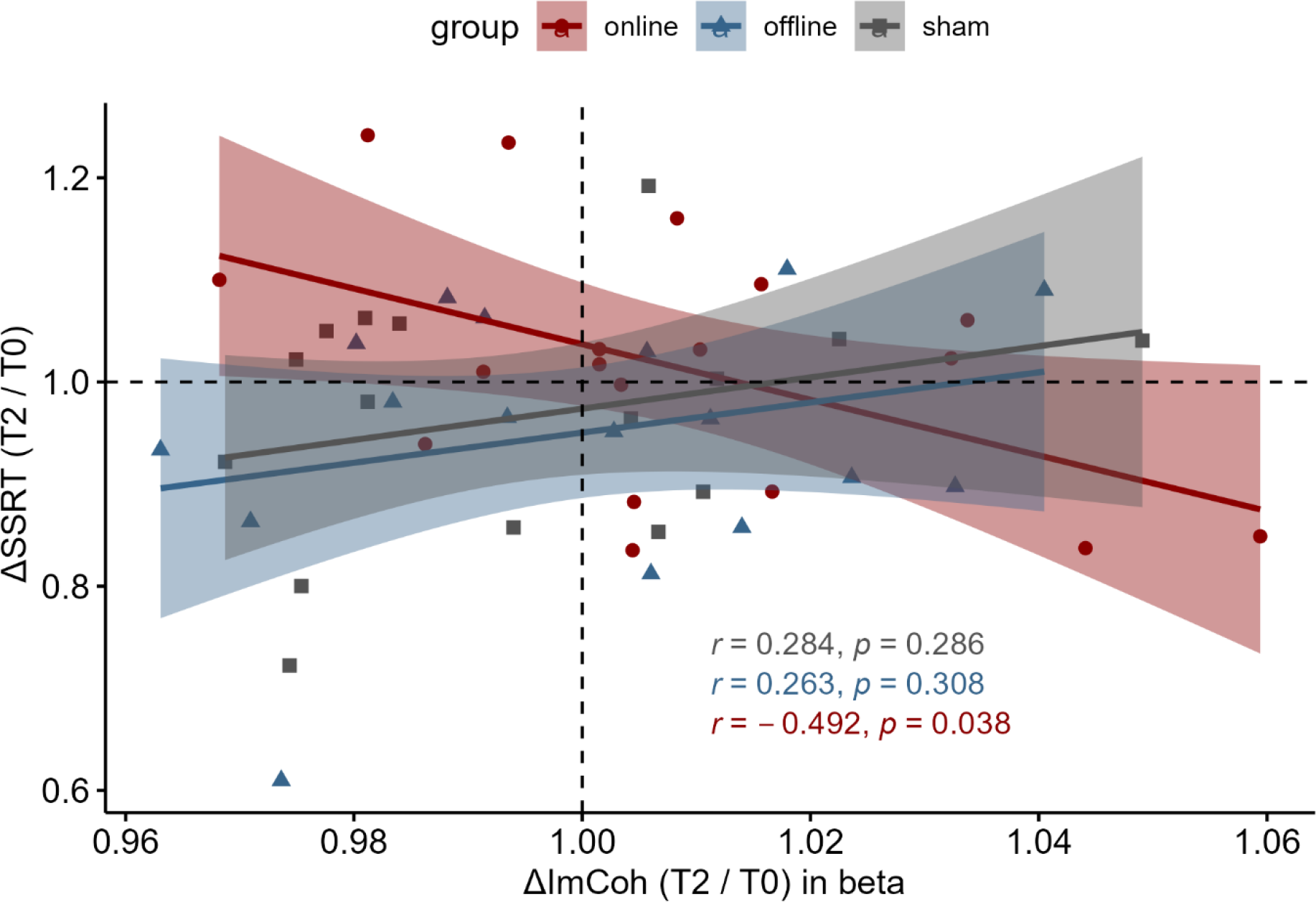
Scatter plot illustrating the relationship between changes in ImCoh in the beta band (13–30 Hz) and changes in SSRT for online, offline, and sham stimulation groups. All p-values are uncorrected.

**Table 3.** Pearson’s product-moment correlation coefficients (r) and associated probability levels (p) (uncorrected) between changes in ImCoh and changes in SSRT from T0 to T2 for each frequency band for each stimulation session

|  | online | offline | sham |
| --- | --- | --- | --- |
| *resting state* |  |  |  |
| 20 Hz | -0.235 | 0.102 | 0.033 |
| alpha (8–12 Hz) | -0.017 | -0.215 | -0.069 |
| beta (13–30 Hz) | **-0.492** * | 0.263 | 0.284 |
| gamma (31–45 Hz) | 0.315 | -0.003 | 0.041 |
| *during successful stop* |  |  |  |
| 20 Hz | -0.289 | -0.007 | -0.214 |
| alpha (8–12 Hz) | 0.224 | -0.022 | -0.468 |
| beta (13–30 Hz) | -0.204 | 0.021 | -0.321 |
| gamma (31–45 Hz) | 0.062 | 0.195 | 0.001 |

\* uncorrected $p < 0.05$

## Discussion

The present study directly compared the efficacy of the online and offline dual-site tACS on functional connectivity between the rIFG and preSMA and associated response inhibition performance. As predicted, the application of dual-site tACS during task performance (i.e., online) significantly improved task-related functional connectivity (i.e., during successful stops) at the target frequency of 20 Hz and the beta band as a whole, supporting the entrainment hypothesis, whereas the offline application did not result in statistically significant changes in ImCoh at the target frequency of 20 Hz and showed a significant *decrease* in task-related ImCoh in the beta range. With respect to the impacts of these changes in functional connectivity on behaviour, although at the group level, the online group did not show significant improvement in SSRT, the correlation analyses suggested that greater increases in ImCoh in the beta band at rest following the online tACS application was associated with greater SSRT improvements. This finding suggests that *online* tACS is capable of inducing neurophysiological changes *and* associated behavioural changes - particularly in those individuals who exhibit the most robust increases in functional connectivity. Despite the lack of changes in functional connectivity at the target frequency range, the *offline* group exhibited significant improvement in SSRT at the group level, demonstrating its efficacy in inducing behavioural change. We postulate that this behavioural change is a result of the neuroplastic change, rather than the entrainment-induced change observed for the online tACS.

### Online dual-site beta tACS increases functional connectivity in the beta band

We hypothesised that dual-site tACS applied to the preSMA and rIFG - key cortical regions in the purported stopping network (Aron et al., 2007), would result in greater increases in functional connectivity between the target regions when it was applied during task performance (online) relative to when stimulation was applied in isolation without a task performance (offline). As predicted, we observed an increase in task-related ImCoh at the target frequency of 20 Hz following the online tACS application, whereas the offline tACS application resulted in a decrease in ImCoh in the beta band (see Figure 4). The improved ImCoh was extended to the entire beta band showing increased ImCoh both at rest and within the task, specifically in the successful stop trials. This is in agreement with the findings from the previous studies using dual-site online tACS during task performance (e.g., Helfrich et al., 2015; Strüber et al., 2014), in which increased coherence between stimulation sites was observed following stimulation. The present study thus provides direct evidence that online, but not offline, dual-site tACS improves functional connectivity between stimulated sites, at for at least ∼20 min post-stimulation (i.e., the time-frame of the assessment points in the current study).

Although the neurophysiological mechanisms underlying the effect of online tACS is still elusive, several potential explanations have been put forward (see Liu et al., 2018 for more details). Of those, stochastic resonance, rhythm resonance, and temporal biasing of spikes (Liu et al., 2018) may potentially explain the effect of online tACS applications on network connectivity and associated response inhibition performance. Stochastic resonance refers to the probability of external stimulation polarising or depolarising neural populations depending on their current activity state (Miniussi et al., 2013), while rhythm resonance refers to the phenomenon when the frequency used for tACS is similar to that of the ongoing endogenous oscillations in the stimulated region. Similarly, the temporal biasing of spikes hypothesis reflects the idea that the spike timing of neurons is regulated by the interaction between the stimulation and endogenous oscillations (Liu et al., 2018). While these potential mechanisms are assumed to affect the spiking of neurons depending on the field strength of the electrical stimulation, the boundaries of mechanisms are not distinct, with several mechanisms simultaneously operating at a given moment (Liu et al., 2018).

In the current study, unlike online tACS, offline tACS did not result in an improvement in functional connectivity between the preSMA and rIFG in the target beta frequency, which suggests that the application of tACS during a task performance may be more beneficial in improving network connectivity, directly influencing temporal changes in brain activity (Veniero et al., 2019).

### Offline dual-site tACS application improves response inhibition despite a decrease in functional connectivity

Although offline tACS application did not improve network connectivity between the preSMA and rIFG, response inhibition performance - which likely involves these brain regions (Aron & Poldrack, 2006) - *was* improved following offline tACS. The observed improvement in response inhibition is in line with previous research reporting improved inhibitory control *after* the application of similar non-invasive brain stimulation (transcranial direct current stimulation) over the preSMA or rIFG (Cai et al., 2016; Ditye et al., 2012; Fujiyama et al., 2022; Jacobson et al., 2011). While the observed correlations between the SSRT improvement and the increase in ImCoh in the beta band at rest following the online tACS was likely a result of improved network connectivity between the preSMA and rIFG, the improved SSRT performance following the offline tACS together with the previous findings (Cai et al., 2016; Ditye et al., 2012; Fujiyama et al., 2022; Jacobson et al., 2011) indicates that a neural plasticity mechanism was involved in the improvements in response inhibition.

The lack of improved functional connectivity between the rIFG and preSMA following the offline tACS can be potentially explained by the timing of the EEG assessment in the current study. For the offline tACS group, post-EEG assessment (T2) was carried out >20 min after the cessation of offline tACS, while post-EEG assessment was immediately after the end of the online tACS. It is possible to speculate that the effect of dual-site tACS on functional connectivity between the preSMA and rIFG was short-lasting (< 20 min) and the effect was abolished for the offline group at T2. Previous offline tACS studies that observed changes in functional connectivity recorded EEG immediately after the cessation of stimulation (e.g., Helfrich et al., 2015; Strüber et al., 2014). There is evidence to suggest that entrainment effects generated by tACS are short-lasting (e.g., ∼ 120 sec; Schwab et al., 2019); therefore, the timing of the rsEEG measurement in the current study might not be optimal for capturing the residual entrainment effects of tACS on oscillatory activity between the preSMA and rIFG. Alternatively, the lack of improved functional connectivity between the rIFG and preSMA following the offline tACS may be explained by the absence of task performance during the stimulation, i.e., brain state-dependent. Nevertheless, future studies may consider including an additional assessment time point in between two stimulations (e.g., between tACS and sham for the offline group) to fully elucidate the effect of online/offline tACS application on functional connectivity between the preSMA and rIFG.

### Effect of dual-site tACS on functional connectivity in broader frequency ranges

The application of 20 Hz dual-site tACS over the preSMA and rIFG not only changed the functional connectivity at the target frequency (i.e., 20 Hz), but it also induced changes in the broader beta range and other frequency ranges. In the offline group, we observed that task-related ImCoh values significantly decreased in beta and gamma, while a significant increase in ImCoh was observed in the alpha band (Figure 4). Similarly, for the online group, ImCoh values at rest significantly increased in the beta but significantly decreased in the alpha band (Figure 3). Previous studies reported that the effects of tACS on brain oscillations extend to other frequency bands than the frequency targeted by tACS (e.g., D’Atri et al., 2019; Nakazono et al., 2020). For example, 5-Hz tACS stimulating fronto-temporal regions for 10 min increased power in the alpha range (8–11.75 Hz) in addition to the targeted theta range (5–7.75 Hz). These results potentially suggest that tACS can induce changes in the activity of other brain regions indirectly, possibly through the stimulation of interconnected networks. The current study extended these previous findings, which reported the EEG power changes by further providing evidence that tACS may induce a global sychronisation effect affecting broader frequency ranges.

### Limitations and future perspectives

In the present study, a predetermined beta (20 Hz) tACS, which appeared to play a functional role in response inhibition (Aron et al., 2016; Swann et al., 2011), was used for all participants. However, it is conceivable that the brain oscillations that underscore successful response inhibition vary between individuals. As such, the use of individually tailored tACS frequency may maximise the efficacy of tACS in improving performance (Veniero et al., 2019; Vosskuhl et al., 2015). This involves identifying a particular oscillation frequency observed in successful task performance using task-related EEG recorded over a cortical network node (Veniero et al., 2019; Vosskuhl et al., 2015). This particular oscillation frequency is then applied by tACS to facilitate task performance. However, this approach is not only time-consuming but was previously reported in a study applying individualised alpha tACS, that the individualised frequency was not a good predictor of whether the entrainment process would be successful (Helfrich et al., 2014). Conversely, Helfrich et al. (2015) reported that when using dual-site tACS at a single frequency of 40 Hz (gamma), if the peak gamma oscillation was closer to 40 Hz, the extent of improvement in coherence between stimulation sites was greater. It would seem that whether individually tailored tACS frequencies are a requisite for optimising improvements in functional connectivity may require further evidence.

Another related issue is that there may not be an obvious individual peak frequency within the beta band for response inhibition. Ali and colleagues (2013) have suggested that although there is an obvious individual alpha frequency in the alpha band, there may not be only one dominant frequency found in other bands. This idea likely explains the observations in previous studies, which reported a range of beta frequencies, i.e., 13-23 Hz, instead of a single frequency, associated with stopping behaviour in response inhibition tasks (Picazio et al., 2014; Swann et al., 2009; Wagner et al., 2018). Similarly, the present study also showed that changes in functional connectivity between the preSMA and rIFG at rest in the entire beta band were associated with the improvement in response inhibition (see Table 3).

Lastly, we observed a mismatch between improvements in functional connectivity and behavioural performance, as the increase in functional connectivity did not translate to a group-level improvement in behaviour for the online group, while the offline group showed improvement in behaviour in the absence of changes in functional connectivity. The discrepancy is not uncommon in the field of non-invasive brain stimulation, e.g., Perini et al., 2020, possibly suggesting that NiBS is capable of inducing changes in behaviour, even in the absence of changes in functional connectivity between different brain regions and changes in functional connectivity does not necessarily translate to behavioural changes. Further research is needed to better understand the mechanisms underlying these changes in functional connectivity and how they relate to changes in behaviour.

## Conclusion

The current study, for the first time, directly compared the effect of the online and offline dual-site tACS on functional connectivity within the response inhibition network. Our results suggested that the application of dual-site tACS targeting the preSMA and rIFG during the stop signal task performance (i.e., online tACS) significantly improved task-related functional connectivity (i.e., during successful stops). In contrast, offline tACS appeared beneficial in improving response inhibition despite the lack of changes in functional connectivity at the target frequency range, suggesting an independent plasticity mechanism to that which drives online tACS effects. These insights have implications for the study design of future tACS research in order to maximise the extent of improvements in functional connectivity and, potentially, associated improvements in cognitive tasks such as response inhibition.

## Acknowledgment

This work was supported by the Neurotrauma Research Program (20193370) awarded to H.F., A. D. T, and J.T. M.R.H. was further supported by the Australian Research Council (ARC) through the Discovery program (FT150100406; DP200101696). We thank Dr Nathan Smith for his contribution to the data collection.

## CRediT authorship contribution statement

**Hakuei Fujiyama**: Conceptualization, Methodology, Formal analysis, Writing - Original Draft, Visualisation. **Alexandra, G. Williams**: Conceptualization, Methodology Investigation, Formal analysis, Writing - Review & Editing. **Jane Tan**: Conceptualization, Methodology, Visualisation, Formal analysis, Writing - Review & Editing, **Oron Levin**: Writing - Review & Editing, **Mark R. Hinder**: Writing - Review & Editing

## Competing interests

The authors report no competing interests.

## Notes

### Competing Interest Statement

The authors have declared no competing interest.

